# Affinity-optimizing variants within the ZRS enhancer disrupt limb development

**DOI:** 10.1101/2022.05.27.493789

**Authors:** Fabian Lim, Genevieve E Ryan, Sophia H Le, Joe J Solvason, Paige Steffen, Emma K Farley

## Abstract

**Summary:** An emerging regulatory principle governing enhancers is the use of suboptimal affinity binding sites to encode tissue-specific gene expression. Here we investigate if optimizing single-nucleotide variants that violate this principle can disrupt tissue-specific gene expression and development. The ZRS enhancer mediates expression of *Shh* in the posterior of the developing limb buds and is critical for limb and digit development. We find that the ZRS contains suboptimal-affinity ETS binding sites. Two human mutations and a synthetic mutation that optimize the affinity of the ETS-A site from 0.15 to 0.25 relative binding affinity cause polydactyly with the same penetrance and severity. Further increasing the affinity of the ETS-A site results in more penetrant and severe phenotypes. The prevalent use of suboptimal affinity binding sites within enhancers to encode tissue-specificity creates a vulnerability within genomes whereby variants that optimize affinity, even subtly, can be pathogenic. This provides a generalizable approach to identify causal variants that underlie enhanceropathies.

**In Brief:** Subtle increases in low-affinity sites underlie human limb defects, while greater increases in affinity lead to more severe and penetrant phenotypes.

**Highlights:** Prediction and validation of pathogenic enhancer variants

Very subtle increases in affinity of low-affinity sites are pathogenic

Penetrance and severity of phenotype scales with increase in affinity

## Introduction

The human genome contains millions of enhancers (Meuleman et al., 2020). These segments of the DNA act as switches to regulate where and when the approximately 20,000 genes are expressed. As such enhancers encode the instructions for tissue-specific gene expression and thus successful development, adult homeostasis, and cellular integrity (Levine, 2010; Long et al., 2016). For example, in the developing limb, the zone of polarizing activity regulatory sequence (ZRS) enhancer ensures precise expression of Sonic hedgehog (*Shh)* in the posterior limb bud which is required for formation of five digits on our hands and feet (Lettice et al., 2003). Single nucleotide changes within enhancers can lead to changes in gene expression that cause developmental defects, evolutionary changes and disease. For example, single nucleotide variants (SNVs) within the ZRS limb enhancer alter digit and limb development (Kvon et al., 2020; Lettice, 2003; Sagai et al., 2004). A SNV within an enhancer for the membrane protein Duffy is associated with malarial resistance (Tournamille et al., 1995). While a SNV within an IRX3 enhancer results in a predisposition to obesity, and a SNV within the DAAM1 enhancer is associated with increased cell migration in several cancers (Smemo et al., 2014; Zhang et al., 2018).

Indeed, genomic analysis suggest that the majority of variants associated with phenotypic variation and disease are located within enhancers, yet we struggle to pinpoint causal variants as they are often embedded within a sea of inert variants (Maurano et al., 2012; Sakabe et al., 2012; Tak and Farnham, 2015; VanderMeer and Ahituv, 2011; Visel et al., 2009). Identifying the causal variants underlying these changes would enable better diagnosis and stratification of patients for different treatment strategies. Furthermore, identifying the mechanistic cause of enhanceropathies can enable novel drug development and treatments. It is therefore critical that we identify the causal variants underlying enhanceropathies and their mechanism. Yet, pinpointing which variants within an enhancer contribute to disease is a huge challenge (Gallagher and Chen-Plotkin, 2018); it is not experimentally nor financially feasible to test all variants alone or in combinations in every disease and tissue to find the causal enhancer variants within each individual. Instead, we need mechanistic and generalizable principles that can help predict causal variants and then test these predictions. Such an approach would enable systematic and scalable approaches that harness the full potential of genomic data to better human health.

Enhancers control gene expression by binding transcription factors to sequences within the enhancer known as binding sites (Anguita et al., 2004; Reddy and Shen, 1991; Zhou et al., 1999). Enhancers typically bind several different transcription factors that work together to mediate combinatorial control of gene expression (Heinz et al., 2010; Liu and Posakony, 2012; Small et al., 1992; Spitz and Furlong, 2012; Swanson et al., 2010). An emerging regulatory principle governing enhancers is the use of suboptimal or low-affinity binding sites to encode tissue-specific gene expression (Crocker et al., 2016; Datta et al., 2018; Farley et al., 2015b, 2016; Jiang and Levine, 1993; Ramos and Barolo, 2013; Rowan et al., 2010). The use of low-affinity sites for transcriptional activators ensures the enhancer is only active in cells where the combination of factors is present at the right concentrations. Changing all binding sites for a transcriptional activator within an enhancer from low to high affinity leads to loss of tissue-specific expression and combinatorial control (Farley et al., 2015b). The modified enhancer drives expression in all cell types where even low concentrations of the individual activators are expressed (Crocker et al., 2015; Farley et al., 2015b). Based on these previous findings we hypothesized that SNVs that increase the binding affinity of a single site for a transcriptional activator could violate this regulatory principle leading to gain of function (GOF) enhancer activity, ectopic expression and changes in organismal-level phenotypes.

To investigate the hypothesis that affinity-optimizing SNVs disrupt development, we focus on one of the most heavily studied vertebrate enhancers, the ZRS enhancer. This enhancer regulates expression of *Shh* in the posterior margins of the developing forelimb and hindlimb buds in a region known as the zone of polarizing activity (ZPA) and is critical for limb and digit development in chick, mouse and humans (Riddle et al., 1993; Saunders, 1968; Williamson et al., 2016). This approximately 800bp enhancer is highly conserved in sequence between mouse and human, and in both species it is located nearly 1MB away from the *Shh* promoter (Lettice et al., 2003, 2002; Sagai et al., 2005).

To date, 29 human SNVs within the ZRS have been associated with polydactyly (extra digits) and other limb defects such as tibial hemimelia (shortening of the tibia, a long bone of the leg) (reviewed in (Kvon et al., 2020; VanderMeer and Ahituv, 2011)). In other species such as mouse, cat, and chick, SNVs within the ZRS also cause polydactyly and limb defects (Dorshorst et al., 2010; Dunn et al., 2011; Knudsen and Kochhar, 1981; Lettice et al., 2008; Maas et al., 2011; Masuya et al., 2007; Zhao et al., 2009). Indeed, mouse provides an excellent system to study the genetic basis of polydactyly as there is a high degree of conservation in regulation of digit formation between mouse and human. This is exemplified by the fact that SNVs at the same position within the ZRS in both mouse and human cause polydactyly (Masuya et al., 2007; Norbnop et al., 2014). Reporter assays analyzing the impact of SNVs on ZRS enhancer activity in mice suggest that polydactyly is associated with GOF ectopic enhancer activity in the anterior limb bud. Only eight human SNVs associated with polydactyly have been tested within the endogenous mouse locus with four of the eight SNVs leading to changes in digit number (Table S1) (Kvon et al., 2020; Lettice et al., 2017). For the four SNVs that cause polydactyly, GOF enhancer activity occurs, however the underlying genetic mechanisms driving this GOF expression are poorly understood (Kvon et al., 2020; Lettice et al., 2017, 2008). Given the remarkable concentration of variants implicated in human limb defects within the ZRS, the conservation of these phenotypes to mouse, and the ease of phenotyping, the ZRS enhancer provides an ideal system to test our hypothesis that affinity-optimizing SNVs disrupt development.

The ZRS is regulated by a combination of transcription factors including Hand2, HoxD, ETV4/5, ETS-1 and GABPa (Lettice et al., 2017, 2012; Osterwalder et al., 2014; Peluso et al., 2017). Five annotated sites known as ETS1-5 are involved in transcriptional activation of *Shh* from the ZRS and bind the transcription factors ETS-1 and GABPa (Lettice et al., 2012). Both ETS-1 and GAPBa are activated downstream of fibroblast growth factor (FGF) signaling from the apical ectodermal ridge (AER) (Ohuchi et al., 1997). Deletion of these five ETS sites (ETS1-5) results in complete loss of enhancer activity within the ZPA when tested by reporter assays in mice (Lettice et al., 2012). However, deletion of individual sites has no impact on expression, which is thought to be due to redundancy between the five ETS sites (ETS1-5). Consistent with this hypothesis, deleting combinations of these sites leads to significant reduced expression within the ZPA (Lettice et al., 2012). Taken together, these results demonstrate that these five ETS sites (ETS1-5) are critical for activation of *Shh* expression in the ZPA.

Consistent with findings in other developmental enhancers regulated by pleiotropic factors, we discover that ETS1-5 are suboptimal affinity binding sites (Figure 1A). We also identify another ETS site, named ETS-A, which is conserved between mouse and human and has an affinity of 0.15 relative to consensus. Two human SNVs associated with polydactyly reside within this low affinity ETS-A site (Albuisson et al., 2011; Kvon et al., 2020). Strikingly both of these mutations increase the affinity of ETS-A by 10% to a final relative affinity of 0.25 (Figure 1C). We hypothesized that this subtle increase in binding affinity could be driving polydactyly. To test this, we created mice with each of these affinity-optimizing variants and an additional mouse line with a sequence change that is not found in humans but increases the affinity of ETS-A by the same amount. Shockingly, all three ETS-A sites with the same subtle increase in affinity cause polydactyly with the same penetrance and severity (Figures 2, 3, 4). Increasing the affinity of the ETS-A site further to 0.52 relative binding causes more penetrant and severe polydactyly, and defects in the tibia (a long bone in the leg) (Figures 5, 6). In contrast, as expected due to the redundancy of binding sites within the ZRS, the loss of ETS-A has no impact on gene expression or limb development (Figure 3). This work demonstrates that affinity-optimizing variants, even ones that only slightly increase low-affinity binding sites, can drive ectopic expression and developmental defects. Furthermore, this study demonstrates a framework for using regulatory principles to understand enhancers and violations in these principles to pinpoint causal variants underlying enhanceropathies.

**Figure 1:**
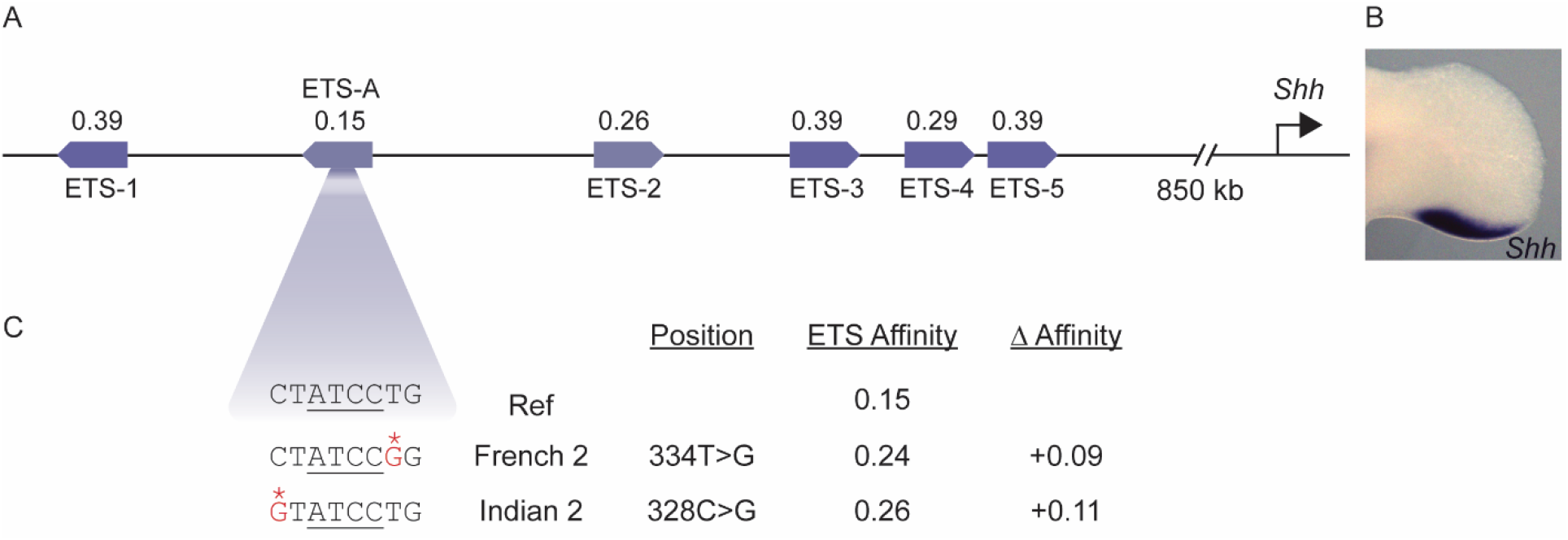
The ZRS contains suboptimal-affinity ETS sites and a newly identified ETS-A that harbors two human variants associated with polydactyly, both human SNVs subtly increase the ETS-A affinity. **A**. The human ZRS contains five annotated ETS sites, ETS1-5 all of which are suboptimal affinity. We also identify a new site, ETS-A, which has a relative affinity of 0.15. **B**. *In situ* hybridization of e11.5 mouse developing hindlimb shows *Shh* expression in the posterior of the limb bud in a region known as the ZPA. **C**. Two human SNVs associated with polydactyly, denoted French 2 and Indian 2, occur within the ETS-A site. The reference ETS-A sequence is shown for comparison. Both SNVs lead to a subtle 10% increase in the affinity of ETS-A to 0.24 and 0.26, respectively.

**Figure 2.**
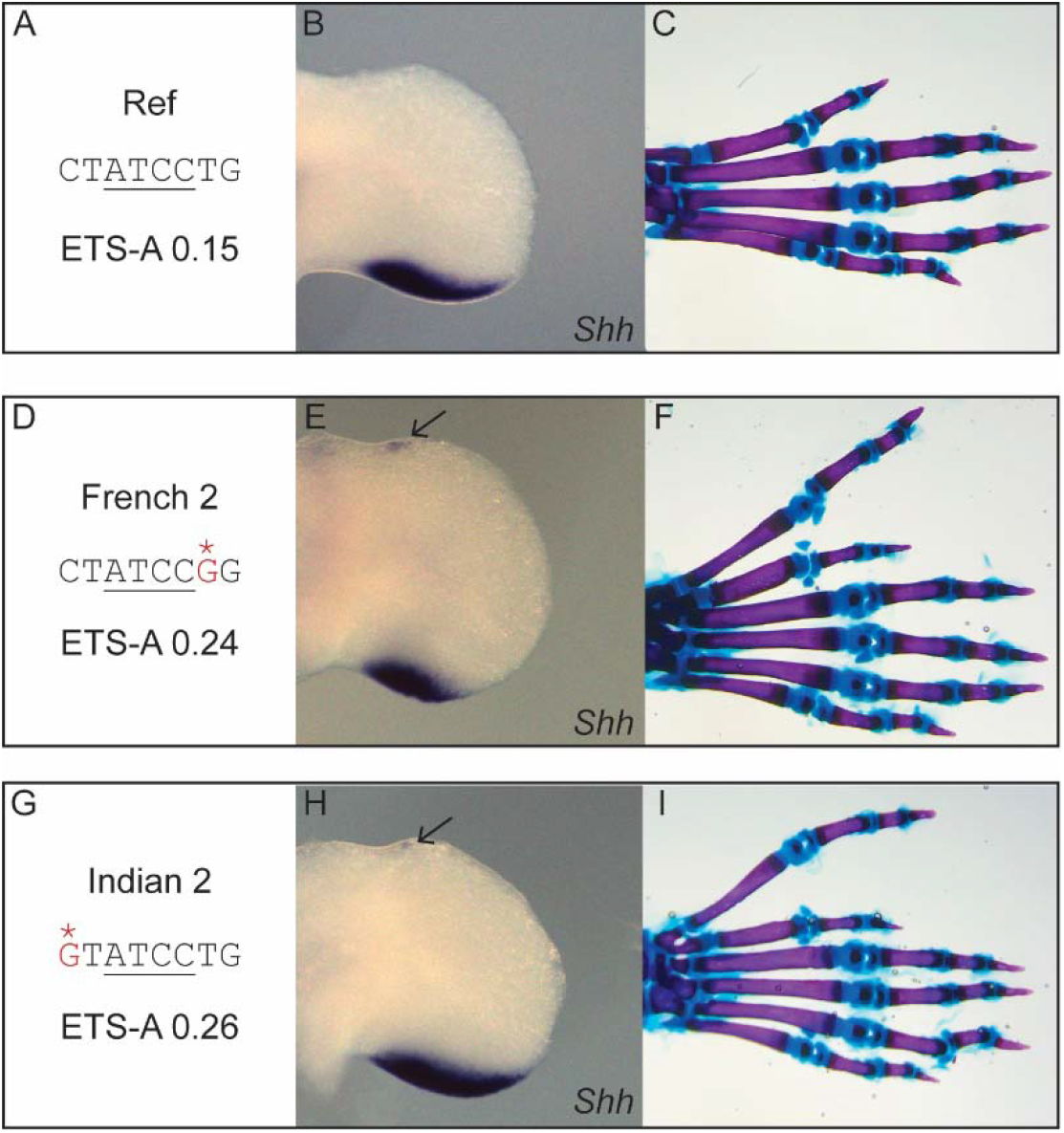
Indian and French 2 SNVs cause polydactyly in mice and phenocopy the human polydactyly phenotypes. **A**. The ETS-A sequence in the reference mouse and human genomes. **B**. *In situ* hybridization of *Shh* in the WT (Ref) developing hindlimb at e11.5 shows a domain of expression in the posterior of the limb bud. **C**. Skeletal staining of a WT hindlimb showing five digits. Digits 2-5 have three bones (are triphalangeal) while the thumb has two bones (biphalangeal). **D**. The ETS-A sequence found in the French 2 family contains a change of T to G. **E**. *In situ* hybridization of *Shh* in the developing hindlimb of a homozygous e11.75 embryo with the French 2 SNV, in addition to the normal domain of posterior expression there is ectopic expression in the anterior of the limb bud as shown by arrow. **F**. Skeletal staining of a homozygous French 2 mouse hindlimb showing six digits, the extra digit is a triphalangeal thumb. **G**. The ETS-A sequence with the Indian 2 C to G SNV. **H**. *In situ* hybridization of *Shh* in the developing hindlimb of a homozygote with the Indian 2 mutation at e11.75, in addition to the normal domain of posterior expression there is ectopic expression in the anterior of the limb bud as shown by arrow. **I**. Skeletal staining of a homozygous Indian 2 mouse hindlimb showing six digits, the extra digit is a triphalangeal thumb.

**Figure 3.**
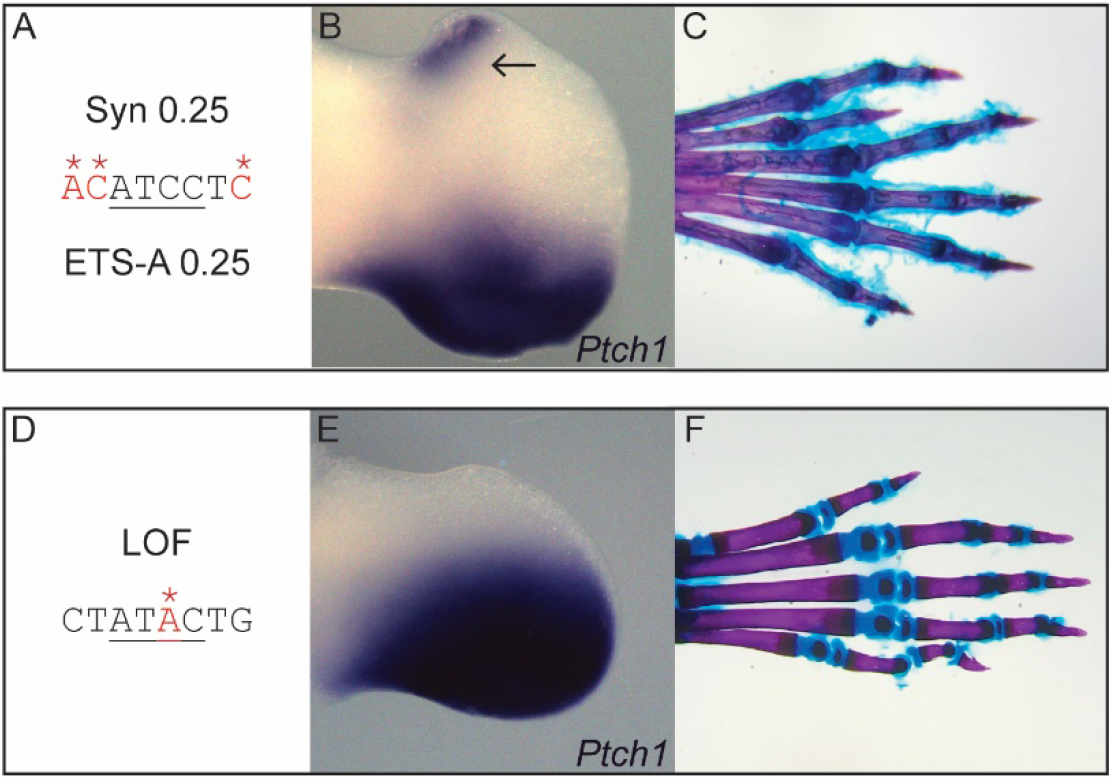
Synthetic changes to ETS-A site creating an ETS-A site with 0.25 affinity and a LOF ETS-A site cause predicted phenotypes. **A**. The Syn 0.25 ETS-A sequence which creates a 0.25 affinity ETS-A site. **B**. *In situ* hybridization of *Ptch1* in the developing hindlimb of a homozygous Syn 0.25 e12.0 embryo, ectopic expression in the anterior of the limb bud as shown by arrow. **C**. Skeletal staining of a Syn 0.25 homozygous mouse hindlimb showing six digits, the extra digit is a triphalangeal thumb. **D**. The ETS-A sequence in the LOF ETS-A site. **E**. *In situ* hybridization of *Ptch1* in the developing hindlimb of a e11.5 LOF embryo shows a domain of expression in the posterior of the limb bud. **F**. Skeletal staining of a homozygous LOF ETS-A mouse hindlimb showing five digits with normal morphology.

**Figure 4.**
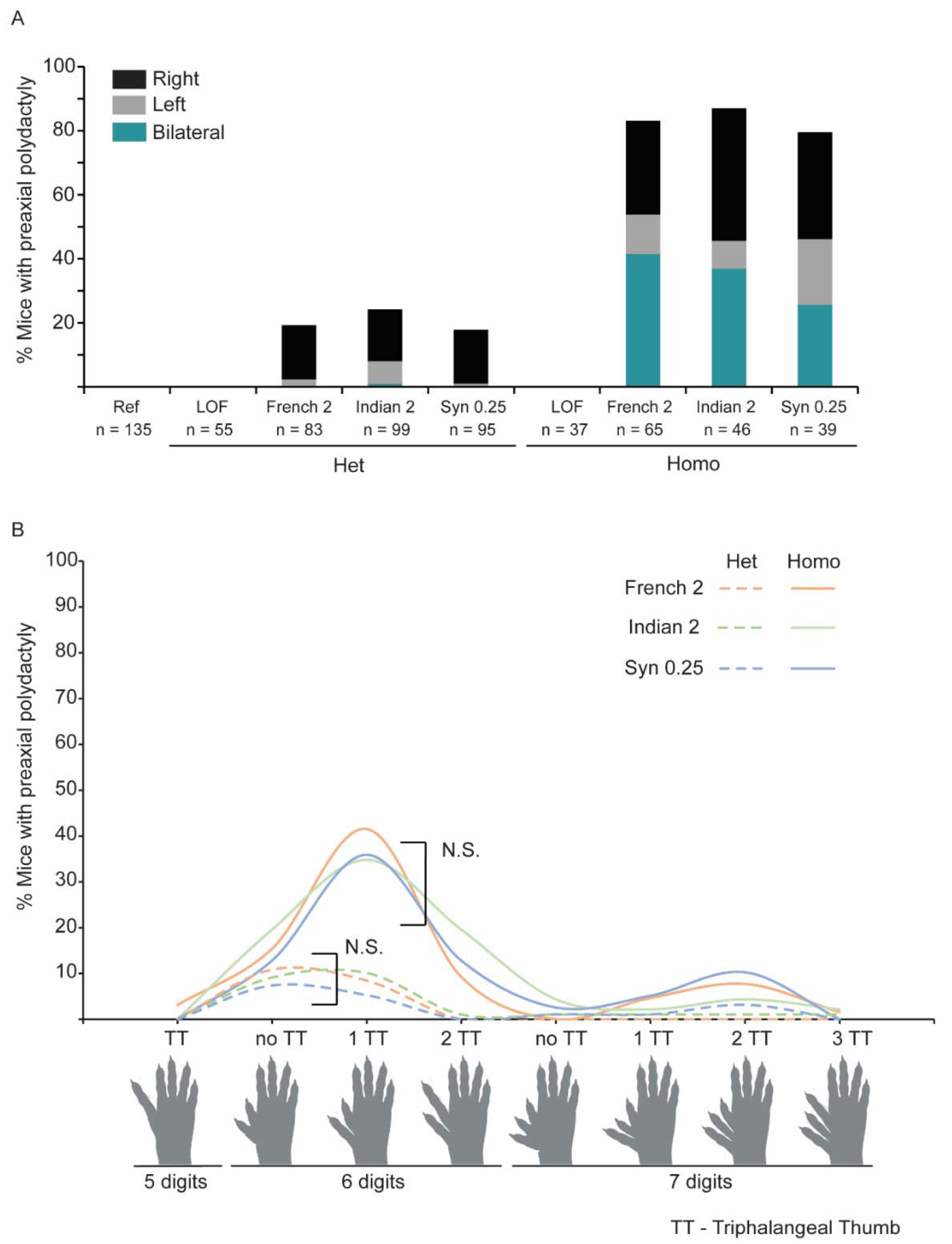
Penetrance and phenotypes for all three mice with the ∼0.25 affinity ETS-A are indistinguishable. **A**. Penetrance and laterality of phenotypes seen in the French2, Indian2 and Syn 0.25 mice in heterozygotes and homozygotes. There is no significant difference in % penetrance and laterality between heterozygotes and between homozygotes. We observe that the polydactyly occurs more frequently on the right hindlimb in both heterozygotes and homozygotes. **B**. Polydactyly phenotypes seen in French 2, Indian 2 and Syn 0.25 heterozygotes (dashed lines) and homozygotes (solid lines). There is no significant difference in the phenotypes seen between the three lines when tested using a 2x9 Fisher’s Exact Test. See Table S2 for details. TT denotes triphalangeal thumb.

**Figure 5.**
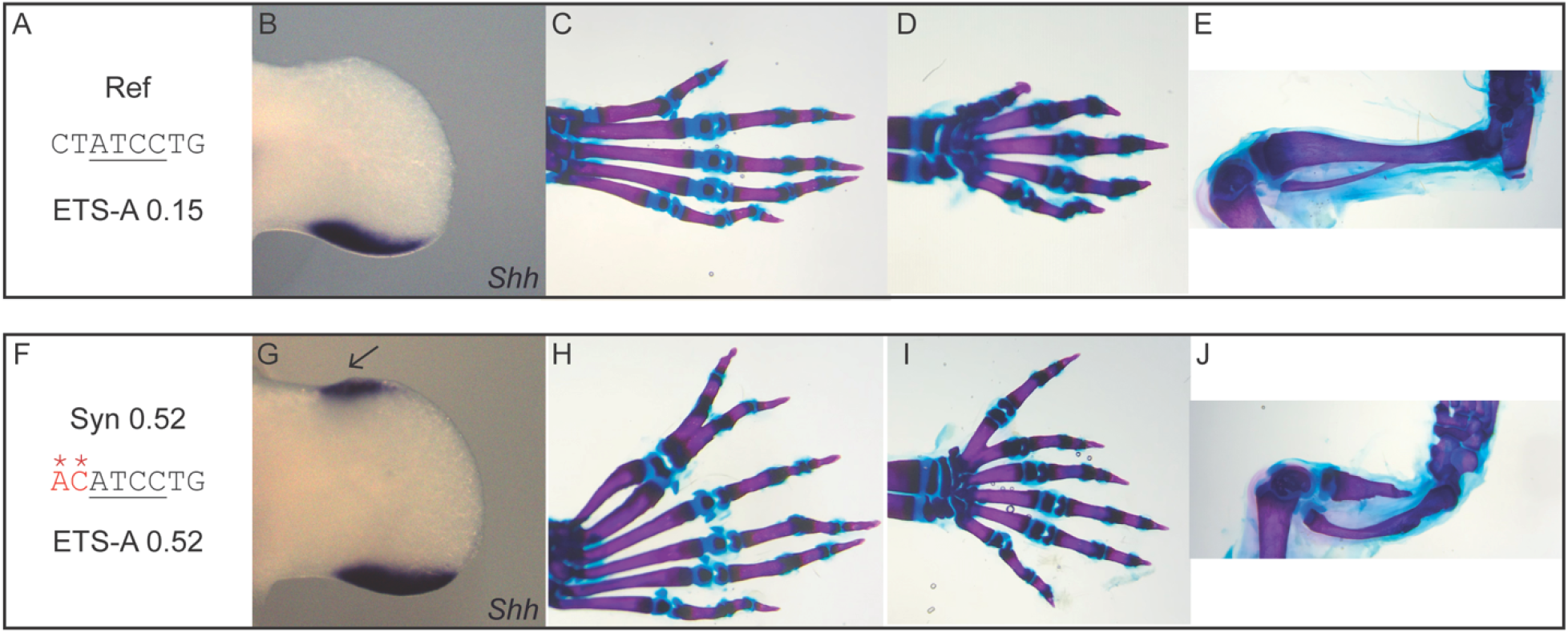
A higher affinity ETS-A site causes polydactyly on forelimbs and hindlimbs and tibial hemimelia. **A**. The ETS-A reference (Ref) sequence. **B**. *In situ* hybridization of *Shh* in the developing hindlimb at e11.5 shows a domain of expression in the posterior of the limb bud. **C**. Skeletal staining of a WT mouse hindlimb showing five digits. **D**. Skeletal staining of a WT mouse forelimb showing five digits. **E**. Skeletal staining of a WT mouse hindlimb focused on tibia and fibula. **F**. The Syn 0.52 ETS-A sequence. **G**. *In situ* hybridization of *Shh* in the developing hindlimb of a e11.75 homozygous embryo with the Syn 0.52 mutation, ectopic expression in the anterior of the limb bud as shown by arrow. **H**. Skeletal staining of a homozygous Syn 0.52 ETS-A mouse hindlimb showing six digits, the thumb and extra digit are both triphalangeal **I**. Skeletal staining of a homozygous Syn 0.52 mouse forelimb showing six digits, both the thumb and extra digit are triphalangeal. **J**. Skeletal staining of a hindlimb focused on tibia and fibula in a Syn 0.52 ETS-A homozygous mouse, the tibia is severely shortened. E & J are same magnification.

**Figure 6.**
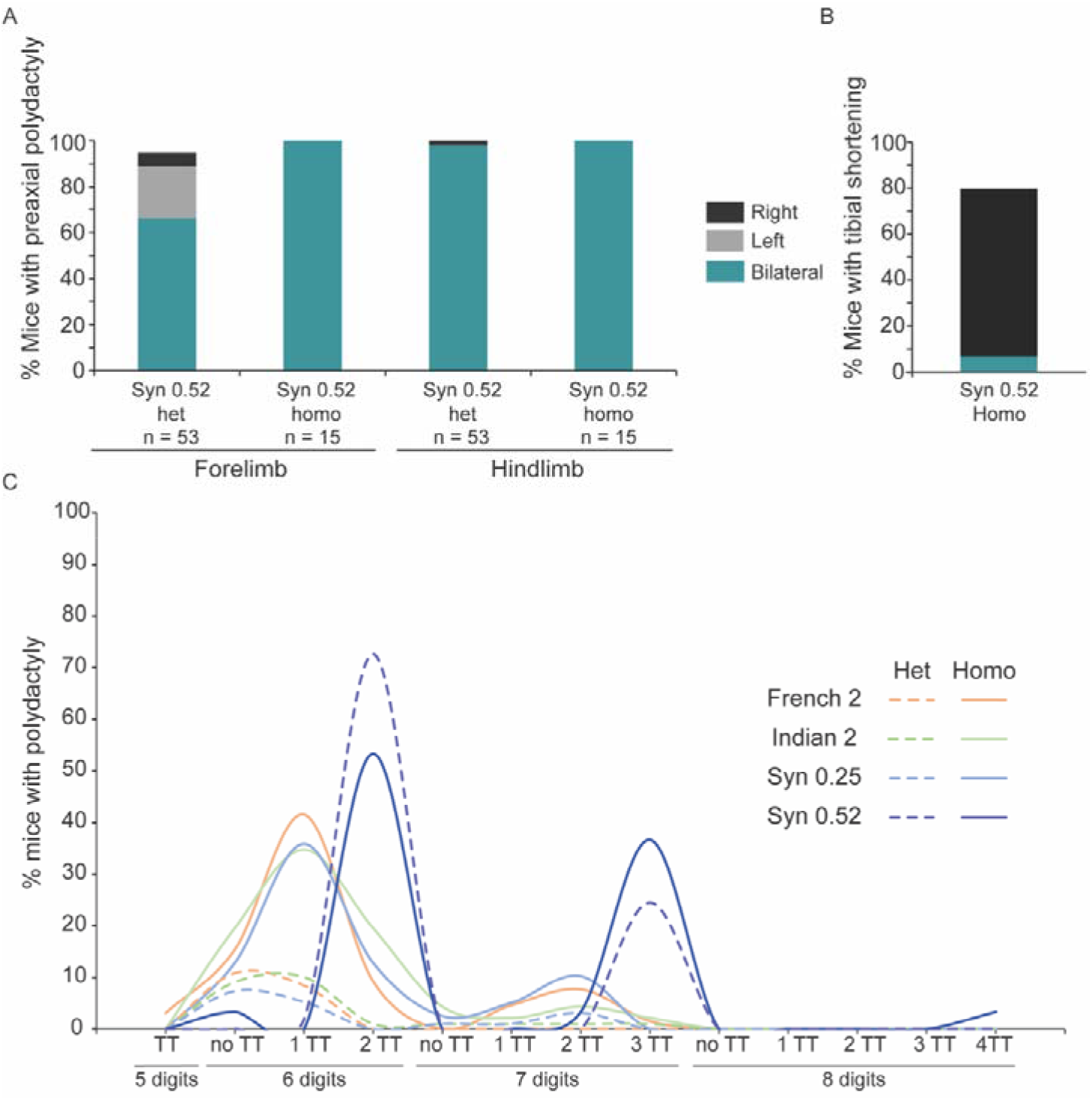
Greater increases in affinity lead to more severe and penetrant phenotypes. **A**. Quantification of penetrance and laterality of phenotypes in both forelimbs and hindlimbs of heterozygous and homozygous Syn 0.52 mice. **B**. Quantification of penetrance and laterality of tibial shortening in homozygous Syn 0.52 mice. No tibial shortening was seen in heterozygous Syn 0.52 mice. **C**. Quantification of polydactyly phenotypes seen in French 2, Indian 2, Syn 0.25 and Syn 0.52 heterozygotes (dashed lines) and homozygotes (solid lines). The Syn 0.52 ETS-A affinity site leads to more severe polydactyly with the most extreme case showing eight digits with four triphalangeal thumbs. See Table S2 for details.

## Results

### The ZRS enhancer is regulated by suboptimal-affinity ETS sites, including the newly identified ETS-A site

An emerging regulatory principle governing enhancers is the use of suboptimal or low-affinity binding sites to encode tissue-specific gene expression. This regulatory principle has been seen in many developmental enhancers including within enhancers regulated by ETS transcription factors (Crocker et al., 2015; Farley et al., 2015b, 2016). To investigate if the ZRS also adheres to this regulatory principle, we measured the relative affinity of these five sites ETS sites (ETS1-5) using Protein Binding Microarray (PBM) data for mouse ETS-1 (Wei et al., 2010). PBM measures the affinity of all possible 8-mers for the transcription factor of interest to provide a direct measurement for every 8-mer sequence. A relative affinity is then calculated by comparing the signal of all 8-mers to the signal of the highest affinity site (Bulyk, 2007; Hume et al., 2015). The 8-mer sequence that binds ETS the strongest has a score of 1.00 or 100%, and this sequence is CCGGAAGT in all ETS Class I transcription factors in both mouse and humans (Badis et al., 2009; Wei et al., 2010). Indeed, the DNA binding domain and specificity of ETS-1 and other Class I ETS transcription factors are conserved from flies to humans (Figure S1 and (Nitta et al., 2015; Wei et al., 2010)). Therefore, the binding affinities measured for ETS-1 also convey the binding affinity for other Class I ETS-1 transcription factors that are also expressed within the limb bud and may also bind to this locus such as GABPa (Lettice et al., 2012; Peluso et al., 2017; Ristevski et al., 2004; Wei et al., 2010). Strikingly we find that the five annotated sites have suboptimal binding affinity, ranging from 0.26 to 0.39 affinity relative to consensus (Figure 1A) (Lettice et al., 2012). Searching the ZRS using PBM datasets identified a total of nineteen putative ETS sites within the human ZRS, seventeen of these sites are conserved in location and affinity between human and mouse. One of these conserved sites is a newly identified site called ETS-A which has an affinity of 0.15 (Figures 1A, S2).

### The two human variants cause similar subtle increases in ETS-A binding affinity

The ETS-A site lies within a region of the ZRS that is completely conserved between mouse and human (Figure S2). We noticed that two human variants associated with polydactyly lie within the ETS-A site, these SNVs are known as the French 2 and Indian 2 variants (Figure 1C). The French 2 variant is found in a family; three family members with this variant have an extra thumb, and a fourth family member has the variant but no polydactyly, demonstrating the variant is not fully penetrant (Albuisson et al., 2011). Only one individual with the Indian 2 variant has been identified, this individual has triphalangeal thumb and preaxial polydactyly (Kvon et al., 2020). Notably, both of these human variants cause a similar subtle increase in the relative affinity of the ETS-A site from 0.15 in the reference to 0.24 in French 2 and 0.26 in Indian 2. We wondered if this slight ∼10% increase in the relative affinity of the ETS-A site could be causing polydactyly. While the French 2 and Indian 2 variants have been studied using LacZ reporter assays (Kvon et al., 2020), neither of these human variants have been studied within the endogenous ZRS locus. Therefore, we first sought to determine if CRISPR/Cas9-mediated human variant knock-in mice with the French 2 and Indian 2 variants exhibit ectopic expression of *Shh* and preaxial polydactyly.

Using CRISPR/Cas9, we created two mouse lines each one with the single nucleotide change within the ZRS, one harboring the French 2 T to G variant (French 2) and another with the Indian 2 C to G variant (Indian 2) (Figure 2). French 2 and Indian 2 homozygous mice showed ectopic expression of *Shh* in the anterior of the hindlimb at e11.75 (Figures 2E, H). The domain of ectopic *Shh* is small (Figure 2H) and therefore for the Indian 2 variant, we also looked at *Ptch1* which is a direct target of *Shh* and commonly used as a readout for *Shh* signaling. *Ptch1* is ectopically expressed in the Indian 2 e12.0 homozygotes (Figure S3C). Having seen that the affinity-optimizing variants lead to ectopic expression of *Shh* in the anterior limb bud, we next investigated if this ectopic expression affected limb development.

Both heterozygous and homozygous French 2 and Indian 2 mice have preaxial polydactyly. In both French 2 and Indian 2 heterozygotes, the most common presentation of preaxial polydactyly is an extra thumb that is biphalangeal or triphalangeal, and this phenotype occurs most commonly on the right hindlimb only. In both French 2 and Indian 2 homozygotes, the most common presentation of preaxial polydactyly is an extra triphalangeal thumb, which occurs on both hindlimbs, or on the right hindlimb if unilateral (Figures 2F, I). Thus, both of these variants are causal for polydactyly and phenocopy the observed human phenotype of preaxial polydactyly with an extra thumb (Albuisson et al., 2011; Kvon et al., 2020).

### Affinity change and not sequence drives the phenotypic change

As both variants give the same phenotypes and increase the affinity of ETS-A by the same amount, this indicates that the mechanism driving polydactyly in both of these human mutations could be the very subtle increase in affinity of the ETS-A site. To test this prediction, we made two more mouse lines with manipulations within the ETS-A site. The first mouse line contains a 0.25 ETS-A affinity site, the same affinity as the Indian 2 and French 2 but with a different sequence change (Figure 3A). This mouse line is called Synthetic 0.25 (Syn 0.25), as we synthetically created an ETS-A site of 0.25 affinity. If the affinity change, rather than the specific sequence change, is driving the phenotype then the Syn 0.25 mice should have the same phenotype as French 2 and Indian 2 mice.

It is possible that any disruption to the ETS-A sequence could lead to a phenotype. To demonstrate that this is not the case we also made a mouse line that we predicted would have no impact on phenotype (Figure 3D). Reporter assays to study the role of the five ETS sites (ETS1-5) find redundancy, such that a reduction in expression is only seen when combinations of these sites are deleted (Lettice et al., 2012). This is consistent with other studies on enhancers that find redundancy of transcription factor binding site within enhancers (Spivakov, 2014). Based on these findings we predicted that the loss of ETS-A site would have no impact on *Shh* expression or limb development. We created an ETS-A loss of function (LOF) mouse that ablates the ETS-A binding sites by removing a critical nucleotide required for binding (Lamber et al., 2008; Wasylyk et al., 1991).

Mice harboring the Syn 0.25 ETS-A site show ectopic expression of *Ptch1* in the anterior limb bud at e12.0 (Figure 3B). In Syn 0.25 heterozygotes, the most common presentation of preaxial polydactyly is an extra thumb that is biphalangeal or triphalangeal. This phenotype occurs most commonly on the right hindlimb. In Syn 0.25 homozygotes the most common presentation of preaxial polydactyly is an extra triphalangeal thumb, occurring most commonly on both hindlimbs, or on the right hindlimb if unilateral (Figure 3C). Thus, the Syn 0.25 ETS-A mice phenocopy the French2 and Indian2 mice. In contrast heterozygote and homozygote mice for the LOF mutation show no ectopic expression (Figure 3E) and all mice have normal limb morphology (Figure 3F). Together these studies demonstrate that the GOF increase in ETS-A affinity within the ZRS enhancer is pathogenic while the LOF variant is non-pathogenic.

### Quantifying phenotypes: All three mutations that increase the affinity by ∼10% lead to the same penetrance, laterality and severity of polydactyly

While qualitatively our prediction regarding the role of affinity-optimizing variants within ETS-A site appears to be accurate, if all three of these ETS-A affinity-optimizing variants are the same then the penetrance, laterality and severity of polydactyly should be comparable between the three lines. We quantified the phenotypes for the Indian 2, French 2 and Syn 0.25 affinity ETS-A mice and compared these to the LOF and reference mice. Phenotyping of mice was done blind to genotype. All three mouse lines with the same affinity ETS-A site have similar penetrance and laterality in heterozygotes with polydactyly occurring unilaterally and most frequently on the right hindlimb (Figure 4A). French 2, Indian 2 and Syn 0.25 homozygotes have a higher penetrance of polydactyly than heterozygotes, and homozygous mice display phenotypes bilaterally. Indeed, in both heterozygotes and homozygotes, all three lines with the same increase in affinity show the same penetrance and laterality with no statistical difference between the three lines (Table S2). This provides further support that the same mechanism, namely a very subtle affinity optimization, is driving the phenotype.

We next looked at the type of preaxial polydactyly occurring within these three mouse lines with 0.25 affinity ETS-A sites (Figure 4B). In the heterozygotes for all three lines preaxial polydactyly with an extra thumb that is either biphalangeal or triphalangeal is most common, while in homozygotes an extra triphalangeal thumb is most common (Table S2). Again, there was no significant difference between the polydactyly phenotypes seen in the three lines with the same affinity change (Table S2). Syndactyly of digits was also observed, this was typically syndactyly of the thumbs and was more prevalent in homozygotes than in heterozygotes. We did not observe any significant difference in phenotypes between male and female mice (Table S2).

This comprehensive analysis demonstrates that all three mutations that increase the affinity of ETS-A to 0.25 have indistinguishable phenotypes in both heterozygotes and homozygotes. This quantitative analysis of phenotypes across 587 transgenic mice provides compelling evidence that the affinity-optimization of this ETS-A site rather than the exact sequence changes is driving polydactyly (Table S2). It is shocking that such subtle affinity-optimization can disrupt development and create extra digits. Furthermore, while there is increasing recognition of the role of low-affinity sites within enhancers, low-affinity sites of such suboptimal affinity as 0.25 are still typically ignored (Kvon et al., 2020; Yan et al., 2021). Yet here we see that a 0.25 affinity site is not only functional but is sufficient to disrupt normal limb development, indicating that subtle increases in low-affinity sites can be pathogenic.

### Greater increases in the affinity of the ETS-A site causes more penetrant and severe phenotypes

Having seen that a subtle increase in affinity can cause developmental defects, we wondered if a greater increase in affinity could cause more severe and penetrant phenotypes. If the degree of affinity change could predict penetrance and severity of phenotype this could be a valuable tool for diagnostic and treatment purposes. To test this prediction, we created a mouse line with an ETS-A site of 0.52 affinity (Figure 5F).

While Syn 0.25, French and Indian 2 homozygote mice have a small amount of ectopic expression within the anterior of the hindlimbs, the 0.52 site lead to a large domain of *Shh* expression in the anterior of the hindlimb (Figure 5G). Additionally, we also see ectopic expression of *Shh* in the forelimbs (Figure S4C). Ectopic expression of *Ptch1* also occurs in the anterior forelimbs and hindlimbs (Figures S3, S4). Having seen that the higher affinity site leads to a greater level of ectopic expression in the hindlimb and forelimb we investigated the impact of this expression on limb development.

As expected, the higher affinity site leads to more severe phenotypes with polydactyly occurring on both the forelimbs and hindlimbs of heterozygous and homozygous Syn 0.52 mice (Figures 5H, I). Syndactyly is seen in the Syn 0.52 heterozygous and homozygous mice more frequently than in the heterozygous and homozygous French2, Indian 2 and Syn 0.25 mice. Furthermore, Syn 0.52 homozygous mice have shortened tibia (Figure 5J). The severe limb and digit defects in Syn 0.52 homozygous mice cause them to have difficulty moving. Thus, as predicted, mice with the greater increase in affinity have more penetrant phenotypes that present bilaterally, and the type of polydactyly is also more severe.

In the most extreme case, we see a Syn 0.52 homozygote mouse with eight digits with four triphalangeal thumbs (Figure 6). The hindlimb phenotypes are fully penetrant for both heterozygotes and homozygotes (Figure 6A). The forelimb phenotypes are fully penetrant in the homozygotes, and almost fully penetrant in heterozygotes. Syn 0.52 forelimb phenotypes are mainly bilateral, but if the phenotype occurs unilaterally, it is seen more frequently on the left forelimb (Figure 6A). Tibial shortening is seen in 80% of homozygotes but never in heterozygotes (Figure 6B). This is most often seen on the right, reflecting the human condition in which 72% of patients with tibial hemimelia have shortening on the right limb (Spiegel et al., 2003). We see no difference in phenotypes between male and female mice (Table S2). Comparison of the types of polydactyly seen in the 0.52 ETS-A mice compared to the three lines with a 0.25 affinity ETS-A site show that the 0.52 ETS-A site causes more severe phenotypes: thus, confirming our prediction that higher increases in affinity lead to more penetrant and severe phenotypes (Figure 6C).

## Discussion

The majority variants associated with disease lie within enhancers (Maurano et al., 2012; Sakabe et al., 2012; Tak and Farnham, 2015; VanderMeer and Ahituv, 2011; Visel et al., 2009). Yet we are unable to pinpoint which enhancer variants are causal. Given the number of enhancer variants associated with disease states and phenotypic variation, it is critical that we develop approaches to reliably differentiate between variants with and without phenotypic consequences. Here we use a mechanistic understanding of the rules governing tissue-specific enhancer activity to predict variants that lead to loss of tissue-specific expression and phenotypes (Farley et al., 2015a, 2015b). We have phenotyped over 700 mice across five transgenic lines, French 2, Indian 2, Syn 0.25, LOF and Syn 0.52 and compared these to reference mice to evaluate our prediction that affinity-optimizing mutations disrupt development and drive polydactyly. We demonstrate that subtle increases in a low-affinity binding site are pathogenic. Furthermore, we identify two rare human SNVs that use this mechanism of action. We also find that a greater increase in affinity leads to more penetrant and severe phenotypes.

### Suboptimal affinity sites within the ZRS encode tissue-specific expression of *Shh* within the limb bud

We and others have contributed to the wealth of data that demonstrate that low-affinity binding sites are critical for tissue-specific gene expression of enhancers (Crocker et al., 2015; Farley et al., 2016, 2015b; Jiang and Levine, 1993; Lettice et al., 2012; Tsai et al., 2017). However, the ZRS is different to other developmental enhancers that have been studied. Firstly, while the majority of developmental enhancers appear to be redundant, deletion of the ZRS leads to loss of *Shh* expression, and loss of the distal limbs and digits (Sagai et al., 2005). The enhancer is also located around 1MB from its target promoter which is further than most well characterized enhancers (Lettice et al., 2002). Finally, the ZRS is highly conserved in sequence between human and all other tetrapods including snakes that no longer have limbs or retain only vestigial limbs (Kvon et al., 2016; Leal and Cohn, 2016). Despite these differences the ZRS, like other enhancers regulated by pleiotropic factors, encodes tissue-specific gene expression using low-affinity sites. This demonstrates that the use of suboptimal affinity binding sites to encode tissue-specific expression is a generalizable regulatory principle that transcends categories of enhancers.

### Gain of function, but not loss of function disrupt development

Enhancers are typically highly redundant. This redundancy ensures robustness within an organism (Cannavò et al., 2016; Hong et al., 2008; Kvon et al., 2021). As a result, loss of a single enhancer is typically compensated by the other redundant enhancers (Osterwalder et al., 2018; Perry et al., 2010). This redundancy occurs at multiple levels. There are typically multiple enhancers known as redundant or shadow enhancers regulating the same gene. Thus, loss of a single enhancer often has no impact on embryonic development or cellular integrity. Another layer of enhancer redundancy is the redundancy encoded within enhancers (Spitz and Furlong, 2012; Spivakov, 2014). This type of redundancy is exemplified by the five ETS sites (ETS1-5) within the ZRS, where loss of a single ETS site has no impact on gene expression (Lettice et al., 2012). Both types of enhancer redundancy mean that loss of a single enhancer or loss of a single binding sites for a transcriptional activator is unlikely to impact expression as this loss can be buffered by other enhancers or redundant binding sites within the enhancer. Meanwhile GOF variants are thought to more likely impact gene expression and development as they lead to ectopic expression that is harder to buffer. As a result, it has been proposed that GOF variants within enhancers are more likely to be causal than LOF variants. Our study supports this hypothesis as we find that loss of the ETS-A has no impact on gene expression, similar to findings using reporter assays to study the role of the ETS1-5 sites. Furthermore, we see no changes in limb development in the 92 ETS-A LOF mice phenotyped, while all four GOF mouse lines we generated drive ectopic gene expression and disrupt limb development.

### Subtle increase in the affinity of TF binding sites can be pathogenic

We hypothesized that the use of low-affinity binding sites within the ZRS enhancer to encode tissue-specificity creates a potential vulnerability within genomes whereby SNVs that optimize the affinity can be pathogenic. Indeed, we find two human variants that lead to incredibly subtle increases in affinity of ETS-A site drive polydactyly with no statistical difference in phenotype between these two variants or the Syn 0.25 site with the same affinity (Table S2). It is shocking that such a subtle change in binding affinity to a final affinity that is still incredibly low can be sufficient to cause GOF enhancer activity and major changes in digit morphology. These studies analyzing phenotypes in over 700 mice demonstrate that affinity-optimizing variants, even subtle ones, can be pathogenic.

### Severity and Penetrance of phenotype increases with affinity

To interpret the role of genetic variation in health and disease and use this knowledge for diagnostic and treatment purposes we need to do more than just pinpoint causal enhancer variants. We also need to predict the likely penetrance and severity of such variants. We predicted and functionally validated that a greater increase in affinity causes more severe and penetrant phenotypes. This is likely true for all changes to affinity that occur within the same binding sites as they are all in the same contextual environment, however the severity and penetrance may also depend on the surrounding binding sites. This interplay of the affinity of binding sites and the dependency between binding sites is known as enhancer grammar and is extensively reviewed in (Jindal and Farley, 2021). Indeed, in our complementary paper by Jindal *et al*. we see that some affinity-optimizing variants have greater impact than expected based on affinity alone and this is presumably due to the interplay of these sites with other binding sites within the enhancer, the enhancer grammar. In the future, studies that integrate an understanding of affinity-optimizing SNVs and enhancer grammar will refine our ability to predict severity and penetrance of affinity-optimizing SNVs.

### Searching for affinity-optimizing variants is a generalizable approach to identifying causal enhancer variants

ETS transcription factors act downstream of the FGF pathway and are implicated in development of all tissues in an organism, including mesodermal induction, pigmentation, cell migration, neural and heart specification (Beh et al., 2007; De Moerlooze et al., 2000; Gainous et al., 2015; Kimelman and Kirschner, 1987; Lanner and Rossant, 2010; Martik and Bronner, 2021; Song et al., 2016; Turner and Grose, 2010). Many diseases result from overactivity of the FGF pathway and hyperactivation of ETS, including cancers, congenital heart disease, neural developmental diseases, craniofacial defects, lung and kidney malformations (Grose and Dickson, 2005; Itoh et al., 2016; Jindal et al., 2015; Turner and Grose, 2010; Xie et al., 2020). Therefore, ETS affinity-optimizing variants within enhancers could contribute to many developmental defects and diseases. Beyond ETS, the role of low-affinity binding sites to encode tissue-specific expression has been described for many other transcription factors such as GATA, Ubx and Prep1 in a variety of tissue types and species (Farley et al., 2015b; Rowan et al., 2010; Tsai et al., 2017). Therefore, we anticipate that affinity-optimizing SNVs that cause GOF enhancer activity and phenotypic change will occur in binding sites for transcriptional effectors that act downstream of most signaling pathways and for pleiotropic factors where suboptimal affinity binding sites are required for tissue-specificity.

The central hypothesis in this study and the accompanying heart study by Jindal *et al*. was that if we can find generalizable rules governing enhancers, then violations in these rules that lead to GOF enhancer activity could pinpoint causal enhancer variants that alter phenotypes. Enhancers are often categorized based on their mode of interaction with the promoter, level of sequence conservation, distance from their target promoter, target gene, tissue of activity, or species. However, here we demonstrate that the use of low-affinity sites to encode tissue-specific expression transcends level of sequence conservation, proximity to promoter, tissue type and even species. The two enhancers studied in these complementary papers are vastly different, one enhancer is active in the *Ciona* heart precursor cells (Beh et al., 2007), while this study focuses on the vertebrate ZRS limb enhancer. The *Ciona* heart enhancer is not conserved at the sequence level and is within 3kb of its target promoter, while the ZRS enhancer is a non-redundant enhancer that is highly conserved and located almost 1Mb from its promoter. Despite these differences both enhancers use suboptimal affinity binding sites to encode tissue-specific expression. Additionally, in both contexts, the use of low-affinity sites creates vulnerability within genomes whereby affinity-optimizing variants can drive ectopic GOF expression that disrupt development. These studies demonstrate the conservation of regulatory principles across diverse enhancers and provide a framework for predicting causal variants underlying enhanceropathies.

## Acknowledgements

We would like to thank all members of the Farley lab for helpful discussions. We thank Kim Cooper for providing advice, use of her microscope and help in identifying the ectopic *Shh* expression via *in situ* hybridization. We thank the Moores UCSD Cancer Center Transgenic Mouse Shared Resource at UCSD and in particular, Jun Zhao for creating our transgenic mice. The authors declare no competing interests.

## Author contributions

E.K.F., F.L., G.E.R. designed experiments. F.L., G.E.R., S.L., P.S. conducted experiments. E.K.F. and F.L. wrote the manuscript. J.J.S. conducted bioinformatic analysis. F.L. and S.L. conducted quantitative analysis of phenotypes. All authors were involved in editing the manuscript.

## Methods

### Key resources table

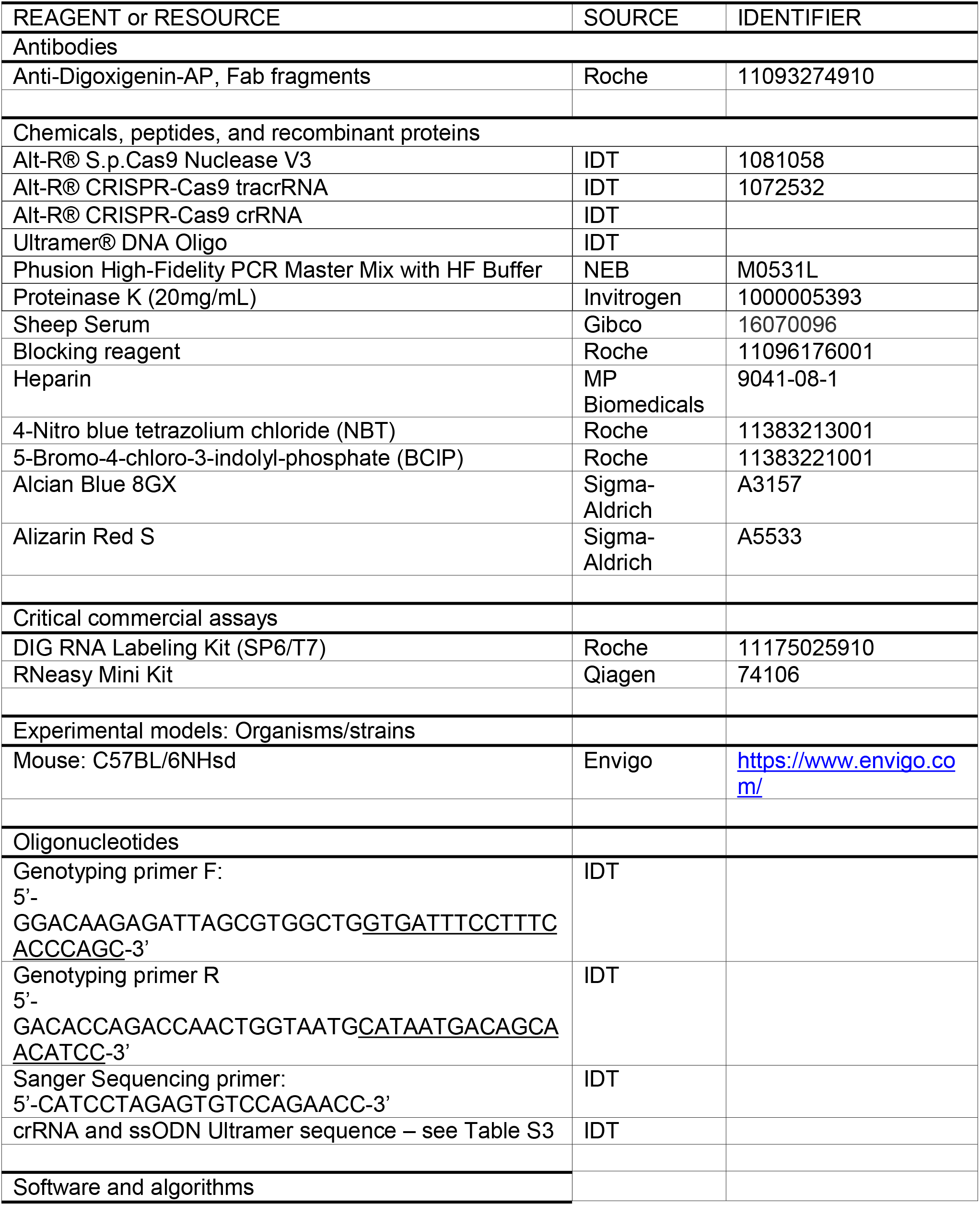
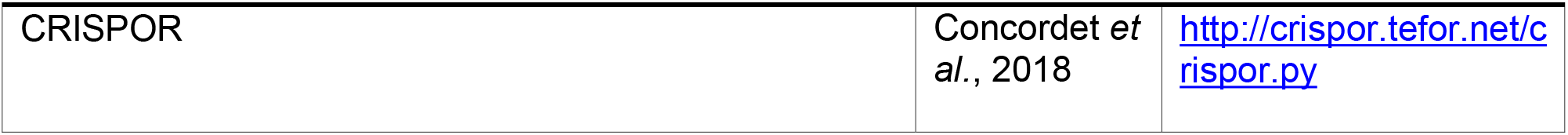

### RESOURCES AVAILABILITY

#### Lead contact

Further information and requests for resources and reagents should be directed to and will be fulfilled by the lead contact, Emma Farley.

### EXPERIMENTAL MODEL AND SUBJECT DETAILS

#### Mice

All experimental procedures were approved by and performed in accordance with the University of California, San Diego Institutional Animal Care and Use Committee. Mice were maintained on a 12:12 light/dark cycle with *ad libitum* standard chow diet and water. Transgenic mouse assays were performed using *Mus musculus* C57BL/6NHsd strain (Envigo). Animals of both sexes were included in this study.

### METHOD DETAILS

#### Generation of transgenic mice using CRISPR-Cas9

Genome-edited mice were generated essentially as described (Yang et al., 2014). Briefly, Cas9 protein, tracrRNA, crRNA, and ssDNA homology-directed repair template oligos were co-injected into one-cell embryos at the Moores UCSD Cancer Center Transgenic Mouse Shared Resource. Custom ssDNA repair oligonucleotides and crRNAs were synthesized by IDT. We designed and selected crRNA if the guide sequence is predicted to have high specificity on CRISPOR (http://crispor.tefor.net/crispor.py) and if the mutation introduced via HDR will ablate the PAM site. Since a PAM site is not available in the genomic locus for the human and synthetic mutations, we first generated a mouse line to contain a *de novo* PAM site within the ETS-A site. French 2, Indian 2, Syn 0.25 and Syn 0.52 mouse lines were generated using CRISPR-Cas9 using one-cell embryos with the new PAM background (see Table S3). LOF mutation mouse line was generated using embryos with the WT background. All mouse lines were generated via homology directed repair using ssDNA as a repair template. Genome-edited founders were identified by genotyping as described below. Wherever possible multiple founders bearing the same desired allele were used to establish each line. Founders were crossed to WT C57BL/6N mice to generate the F1 generation for each mouse line to ensure that potential on-target mutations not detectable by our genotyping assay (e.g., structural variants) were eliminated.

#### Genotyping

Genomic tail DNA was obtained and used to genotype ETS-A transgenic mice with the following primers: Forward 5’-GGACAAGAGATTAGCGTGGCTG<u>GTGATTTCCTTTCACCCAGC</u>-3’ and Reverse 5’-GACACCAGACCAACTGGTAATG<u>CATAATGACAGCAACATCC</u>-3’. The underlined sequences anneal to the ZRS, and the remaining sequences are overhangs used to clone ZRS PCR products into a vector containing an ampicillin resistance cassette by Gibson assembly. For all mice including founders, PCR products were analyzed by Sanger sequencing (sequencing primer: 5’-CATCCTAGAGTGTCCAGAACC-3’) to identify ZRS genotypes. For all founder animals, PCR products were cloned, and individual clones sequenced to confirm initial genotyping results with single allele resolution.

#### Phenotyping

Each mouse born into our colony has all 4 limbs inspected by an investigator blind to genotype at postnatal day 10-18 during routine ear clipping (for identification) and tail biopsy collection (for genotyping). Limb and/or digit phenotypes, or the absence thereof, are readily detectable in postnatal mice and recorded in detail. Specifically, each limb is inspected for number of digits and the presence of other overt features including triphalangeal first digit(s) and/or shortened limbs. Following genotyping, phenotypic data for each genotype within each ETS-A transgenic line is collated to calculate penetrance (based on presence or absence of phenotype).

#### Timed matings for embryo collection

Within each ETS-A transgenic mouse line, timed matings were set up and monitored each morning for vaginal plug formation. The date that plugs were observed was noted as embryonic day 0.5 (e0.5) used to collect embryos at e11.5 and e12.0. Females were removed from males on the plug date and embryos were staged at dissection. Some embryos were identified to be developmentally older than the expected e11.5 embryos and are labeled as e11.75. Pregnant females were humanely euthanized by isoflurane overdose. Embryos were dissected in ice-cold phosphate-buffered saline (PBS) pH 7.4 and then fixed in 4% paraformaldehyde in PBS pH 7.4 overnight with gentle rotation at 4°C. Embryos were then dehydrated through a graded methanol series at 4°C (25%, 50%, 75% MeOH in PBS pH 7.4 plus 0.1% Tween-20, 100% MeOH) and stored in 100% MeOH at -20°C up to 6 months until use. The yolk sac of each embryo was collected and used for genotyping as described above. The sex of embryos is unknown.

#### Probe cloning and synthesis for *in situ* hybridization

*Shh* and *Ptch1* templates were amplified from mouse e11.5 cDNA using primers previously described (Cooper et al., 2014), ligated into a pCR BluntII TOPO vector, transformed into TOP10 competent cells, and plated for selection on kanamycin plates. Colonies were selected for sequence verification and then plasmid prepped. Plasmid DNA was linearized with SpeI or NotI restriction enzyme and then used as a template for *in vitro* transcription using a digoxigenin labeling kit with T7 (antisense) or Sp6 (sense) polymerase. Following DNase treatment to digest template DNA, RNA probes were recovered using a RNeasy mini kit and RNA concentration and purity were measured to confirm probe synthesis.

#### Whole-mount *in situ* hybridization

Embryos were treated with 6% H_2_O_2_ in MeOH for 1 hour, and then rehydrated through a methanol series to PBS-T (1% Tween-20 in PBS pH 7.4). Embryos were washed 5x 5 minutes in PBS-T then treated with proteinase K (10 µg/mL) for 20 minutes. After permeabilization, embryos were washed in PBS-T containing 2 mg/mL glycine, then PBS-T, then post-fixed for 20 minutes in 4% PFA/0.2% glutaraldehyde in PBS-T. Embryos were then washed 2x 5 minutes in PBS-T followed by 10 minutes in a 1:1 mixture of PBS-T and hybridization solution (50% formamide, 5x SSC pH 4.5, 1% SDS, 50 µg/mL yeast tRNA, 50 µg/mL heparin). Embryos were then allowed to sink (no rocking) in hybridization solution for 10 minutes. Embryos were then

changed to new hybridization solution and incubated for at least 1 hour at 65°C. Hybridization solution was replaced with fresh hybridization solution containing 1 µg/mL of antisense (all ETS-A embryos and WT control) or sense (WT control only) probe followed by overnight incubation at 65°C. Embryos were washed 3x 30 minutes in solution I (50% formamide, 5x SSC pH 4.5, 1% SDS) at 65°C followed by 3x 30 minute washes in solution III (50% formamide, 2x SSC pH 4.5) at 65°C. Embryos were then washed 3x 5 minutes in TBS-T (1% Tween-20 in Tris-buffered saline) and blocked for 1 hour in block solution (10% heat-inactivated sheep serum and 0.1% Roche blocking reagent in TBS-T). Roche blocking reagent was dissolved in maleic acid buffer according to manufacturer’s recommendations. Embryos were then incubated in block solution containing 1:2500 anti-digoxigenin-AP antibody overnight at 4°C. Embryos were washed 3x 5 minutes in TBS-T and then 5x 1 hour in TBS-T followed by overnight incubation in TBS-T at 4°C. Embryos were then washed 3x 10 minutes in NTMT (100 mM NaCl, 100 mM Tris pH 9.5, 50 mM MgCl_2_, 1% Tween-20) before coloration in AP reaction mix (125 µg/mL BCIP and 250 µg/mL NBT in NTMT). Coloration was carried to completion in the dark. Embryos were washed 10 minutes in NTMT followed by 3x 10 minutes in TBS-T and then overnight in TBS-T at 4°C. Embryos were imaged using the Leica M165 FC microscope with the Lumenera Infinity3 camera, then post-fixed in 4% PFA for 30 minutes and stored in 1% PFA in 4°C. All steps were performed with gentle rocking and at room temperature unless otherwise specified.

#### Skeletal preparations

Young postnatal mice at age P10-12 were humanely euthanized by CO_2_ inhalation prior to skeletal preparations, with the exception of the Syn 0.25 homozygote at five-months-old. Dissected limbs and/or whole cadavers of representative homozygotes for each line were skinned and eviscerated, then fixed in 95% ethanol overnight. Samples were then stained over two nights in cartilage staining solution (75% ethanol, 20% acetic acid, 0.05% alcian blue 8GX), rinsed overnight in 95% ethanol, cleared overnight in 0.8% KOH, and then stained overnight in bone staining solution (0.005% alizarin red S in 1% KOH). After staining samples were further cleared in 20% glycerol in 1% KOH until digits were free of soft tissue and long bone morphology was visible. Samples were further processed through a graded series of 50% and 80% glycerol in 1% KOH and then into 100% glycerol for imaging and storage. All steps of the skeletal staining procedure were performed with gentle rocking at room temperature.

#### Binding affinity calculation

Relative binding affinity is calculated using high-throughput binding data from the UniProbe database (thebrain.bwh.harvard.edu/uniprobe/index.php). Median intensity signals of 8-mers centering the GGAW core for ETS-1 Protein Binding Microarray data were measured as a percentage of their optimal 8-mer binding motifs (Wei et al., 2010).

## QUANTIFICATION AND STATISTICAL ANALYSIS

To assess any statistical differences in % penetrance and laterality between French 2, Indian 2, Syn 0.25 mice, we performed the Fisher’s Exact Test using the fisher.test function in R. Statistical difference in digit phenotypes were also measured using Fisher’s Exact Test using a 2x9 table. Chi Square Goodness of Fit test was performed to assess if the occurrence of phenotype on the right or left hindlimb among unilateral mice deviate from the assumption that phenotype would occur at a 50%-50% rate on right and left hindlimbs.

## Supplemental Figures

**Figure S1.**
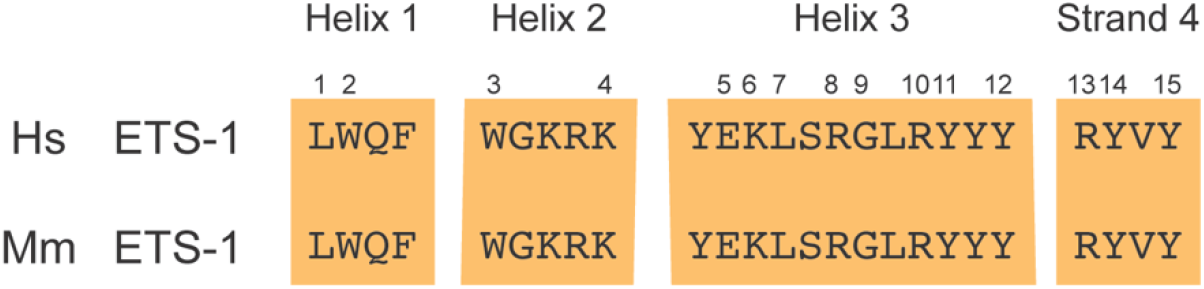
ETS-1 family members have highly conserved DNA binding domain and binding specificity. DNA binding domain of mouse and human ETS I transcription factors. The DNA binding domains are highly conserved. Numbers show the location of contacts between the DNA and protein (Pufall et al., 2005; Wang et al., 2005).

**Figure S2.**
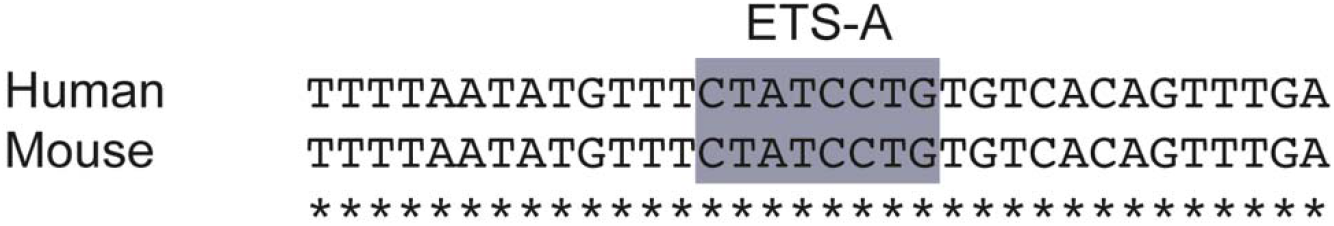
Conservation of ETS-A site between human and mouse. Sequence from human and mouse ZRS around the ETS-A site show perfect conservation, as indicated by asterisks. Blue box highlights the ETS-A sequence within the human and mouse ZRS, both contain a 0.15 affinity ETS-1 binding site as calculated using PBM binding data.

**Figure S3:**
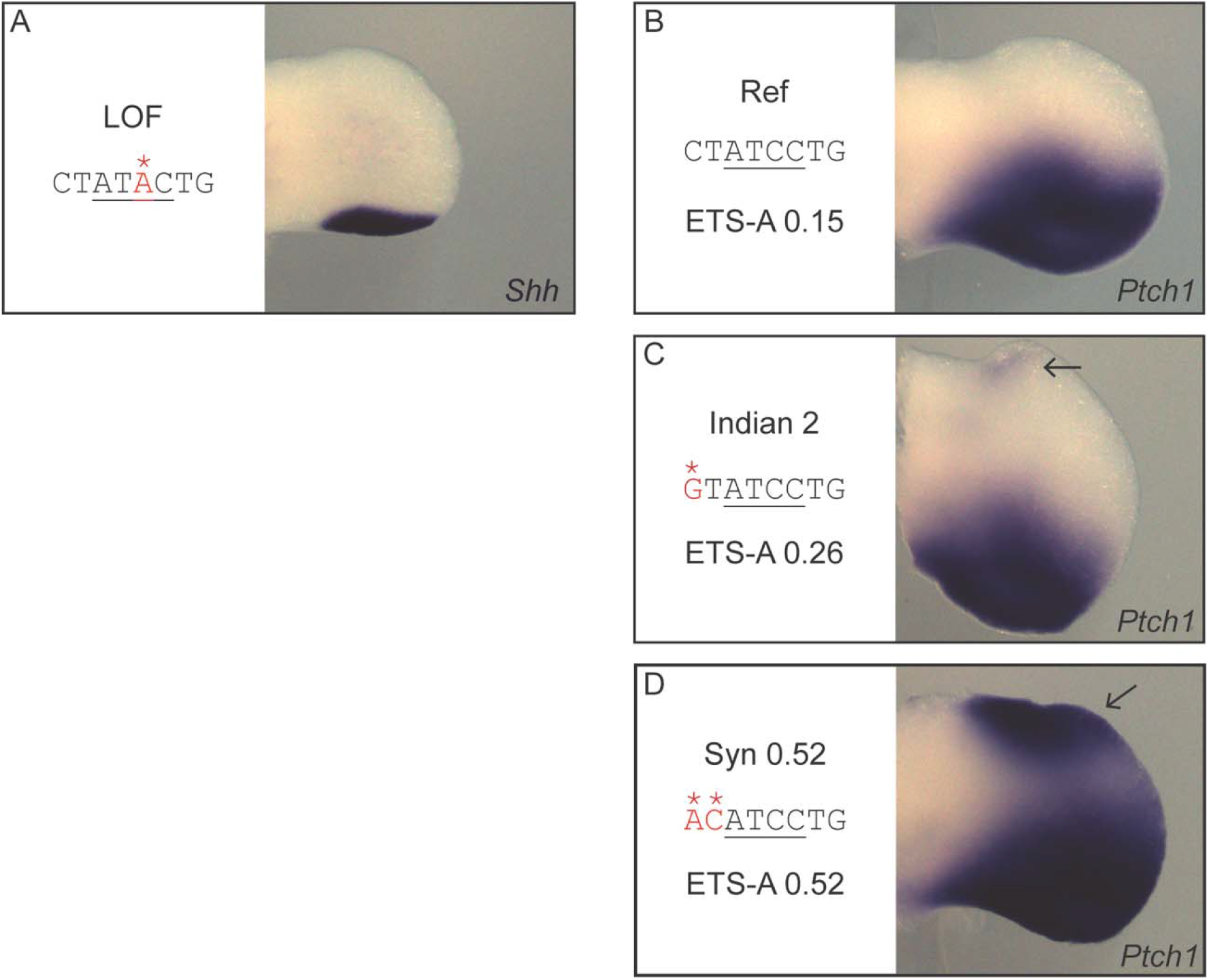
Hindlimb bud *in situ* hybridization of transgenic mice. Embryos were collected at e11.5 to e11.75 for *Shh in situ* hybridization. Embryos were collected at e11.75 to e12.0 for *Ptch1 in situ* hybridization. Ectopic *Ptch1* expression can be seen in Indian 2, Syn 0.52 hindlimb buds. All limb bud images were acquired and cropped using the same settings.

**Figure S4:**
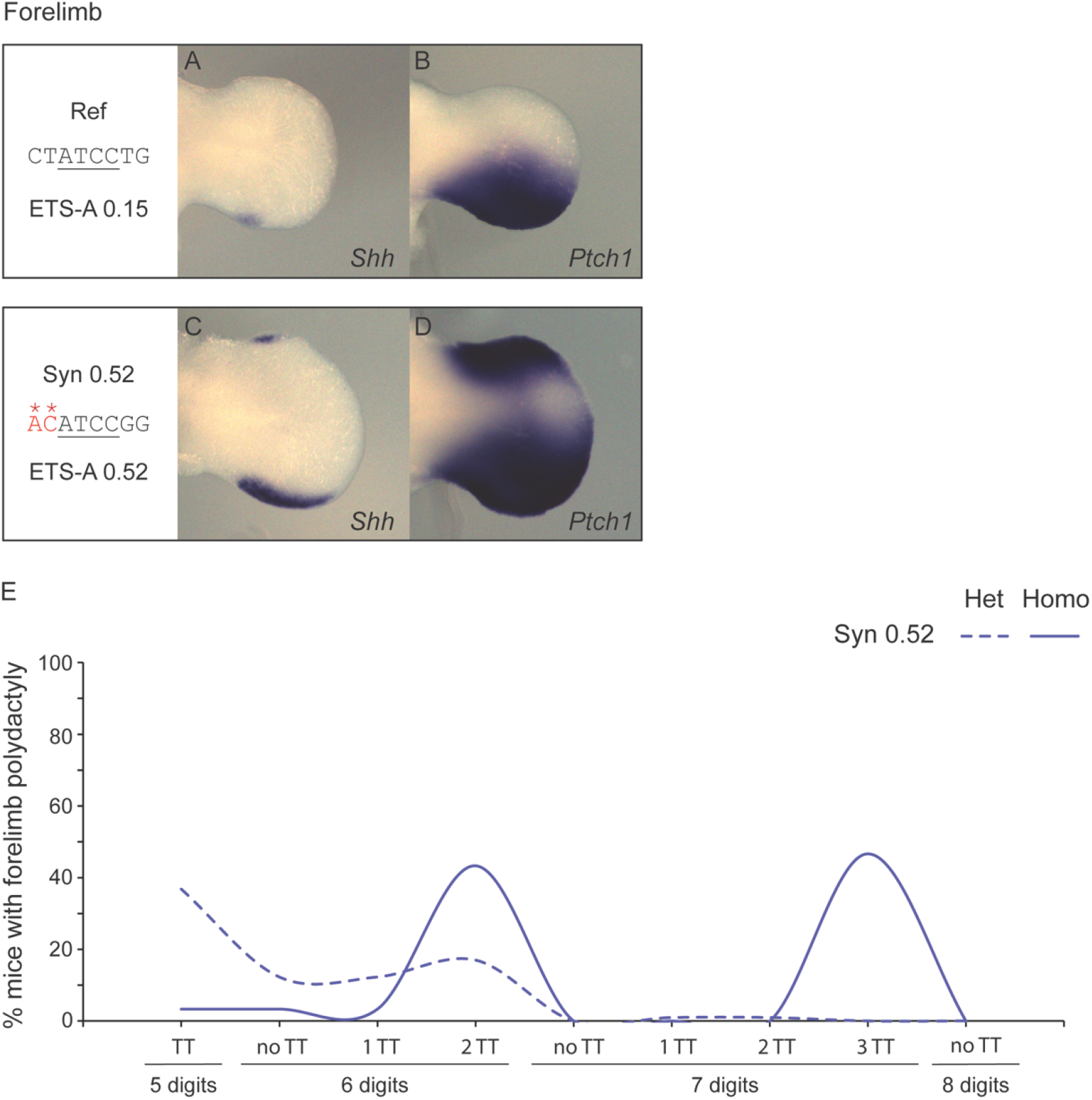
Forelimb phenotypes seen in Syn 0.52 heterozygous and homozygous mice. **A-D.** *Shh in situ* hybridization in e11.5 WT **A & B**, and e11.75 Syn 0.52 embryonic limb buds **C & D**. Ectopic expression can be seen in the anterior forelimb for both *Shh* and *Ptch1*. **E**. Digit phenotypes on the forelimbs of Syn 0.52 mice. Heterozygotes have less severe phenotypes than homozygotes. WT, LOF, French 2, Indian 2 and Syn 0.25 have no forelimb phenotypes.

**Supplementary Table S1**

Description of all human variants tested within the endogenous locus and the phenotypic studies of these mice both in the literature (Kvon et al., 2020; Lettice et al., 2017). and in this study

**Supplementary Table S2**

Phenotypic data of all mice from this study: WT, LOF, French 2, Indian 2, Syn 0.25 and Syn 0.52 mice. **Sheet 1** contains the record of the digit and tibial hemimelia phenotypes from each mouse line, calculated to present the % penetrance and % laterality. **Sheet 2** contains the p-values from Fisher’s Exact Test measuring any significant difference across any pair of mouse lines among French 2, Indian 2, Syn 0.25 in A. % penetrance (two factors: have phenotype or no phenotype) B. % laterality (three factors: bilateral or unilateral or no phenotype) C. Digits phenotype (nine factors: 5 digits no TT, 5 digits 1 TT, ……, 7 digits 3 TT) D. Presentation of digits phenotype occurring to females or males within each mouse line (two groups: e.g. Syn 0.25 HET female vs Syn 0.25 HET male; 9 factors: 5 digits no TT, 5 digits 1 TT, ……, 7 digits 3 TT). **Sheet 3** contains the p-values from Chi Square Goodness of Fit Test used to calculate if the occurrence of phenotype on the right or left limb deviates from the assumption that phenotypes occur on the right and left at a 50%-50% likelihood.

**Supplementary Table S3**

CRISPR/Cas9 components used to create the mice generated in this study. crRNA sequence, genetic background of embryos used for CRISPR injection, and ssODN sequence used to generate each mouse line. All lines were created on a background of C57BL/6NHsd mice.

## Notes

### Competing Interest Statement

The authors have declared no competing interest.

